# JIND: Joint Integration and Discrimination for Automated Single-Cell Annotation

**DOI:** 10.1101/2020.10.06.327601

**Authors:** Mohit Goyal, Guillermo Serrano, Ilan Shomorony, Mikel Hernaez, Idoia Ochoa

## Abstract

Single-cell RNA-seq is a powerful tool in the study of the cellular composition of different tissues and organisms. A key step in the analysis pipeline is the annotation of cell-types based on the expression of specific marker genes. Since manual annotation is labor-intensive and does not scale to large datasets, several methods for automated cell-type annotation have been proposed based on supervised learning. However, these methods generally require feature extraction and batch alignment prior to classification, and their performance may become unreliable in the presence of cell-types with very similar transcriptomic profiles, such as differentiating cells. We propose JIND, a framework for automated cell-type identification based on neural networks that directly learns a low-dimensional representation (latent code) in which cell-types can be reliably determined. To account for batch effects, JIND performs a novel asymmetric alignment in which the transcriptomic profile of unseen cells is mapped onto the previously learned latent space, hence avoiding the need of retraining the model whenever a new dataset becomes available. JIND also learns cell-type-specific confidence thresholds to identify and reject cells that cannot be reliably classified. We show on datasets with and without batch effects that JIND classifies cells more accurately than previously proposed methods while rejecting only a small proportion of cells. Moreover, JIND batch alignment is parallelizable, being more than five or six times faster than Seurat integration. Availability: https://github.com/mohit1997/JIND.

## Introduction

Recent developments in single-cell RNA sequencing (scRNA-seq) technologies have made it possible to profile the transcriptome of thousands of single cells in parallel. Massive amounts of single-cell RNA-seq data can now be generated enabling data-driven studies of gene expression at the single-cell resolution. Applications of this technology include the discovery of new cell-types^1, 2^, identifying potential cellular targets for diseases^3^ and the analysis of cell developmental stages through time^4^.

An important step in single-cell genomic data analysis is the characterization of cell-types in a large mixture of cells. Traditionally, this is done by probing specific marker genes. Thus, a typical pipeline starts with a clustering algorithm to group cells with similar transcriptomic profiles, followed by manual labeling of the clusters based on appropriate biological markers identified in prior studies. However, the variability in clustering methods, the lack of standardized ontologies of cell labels, and the reliance on time-consuming manual annotations make this approach not scalable and creates a bottleneck in single-cell genomics pipelines^5, 6^.

The gain in popularity of single-cell RNA sequencing has led to the creation of very large reference datasets, such as the Human Cell Atlas (HCA)^7^ or the Mouse Cell Atlas (MCA)^8, 9^. These datasets, which are meticulously annotated and extensively validated by researchers, when combined with supervised machine learning techniques, present a natural framework for automating the cumbersome cell annotation process. Based on this idea, several methods have been proposed to transfer labels from an annotated scRNA-seq dataset (source batch) to an unannotated dataset (target batch)^5, 6, 10–15^.

Two questions naturally arise regarding the fundamental limitations of such supervised learning approaches to cell-type identification. First, the source and target batches may exhibit technical variability, generally referred to as *batch effects*, due to differences on data collection or sample preparation. How do these batch effects, which confound true biological differences^16^, affect the reliability of the prediction models that are trained on a source batch and used on a target batch? Second, unlike standard classification tasks where each data point distinctly lies in one and only one class, cells can exist in intermediate states during the process of differentiation^17^. How can these automatic annotation models avoid misclassifying cells that are in transitioning states or cells that are outliers^18^, exhibiting abnormal patterns of gene expression due to the inherent noise in the dataset?

Previously proposed solutions for automated cell identification do not fully address these two fundamental questions. Off-the-shelf classifiers are not well suited for this task as the distribution of the gene expression data can significantly differ between source and target batches. Thus, to handle batch effects, previously proposed solutions either (i) employ classification algorithms that are empirically shown to generalize to datasets with batch effects,^10, 12–14^ or (ii) transform the data in both batches onto a common latent space through dedicated batch alignment methods^15, 19^ (hereafter referred to as *symmetric* alignment methods) prior to training the classifier^11, 15^. On one hand, approach (i) cannot guarantee that the classifier will be robust against arbitrary types of technical variability between the batches. On the other hand, with approach (ii), when new data becomes available, all existing batches must be re-aligned and the prediction model retrained before annotating the new target batch. This significantly increases the computational overhead, and potentially alters previous classification results.

Regarding the classification of cells that lay on intermediate states or at the intersection between different cell-types, previous approaches use a fixed confidence threshold (e.g., 0.9) on the maximum probability across all cell-types, and assign an “unassigned” label to the cells with a lower confidence prediction^10, 11^. However, this does not take into account the variability in the ease of classification across different cell-types, potentially resulting in the filtering (that is, labelling as unassigned) of a large number of cells.

To overcome these issues, we propose a new framework for cell-type identification called JIND. JIND is based on neural networks (NNs) and automatically learns a low-dimensional representation (latent space) from the source batch that is well suited for cell-type classification. To deal with batch effects, JIND projects the target batch onto the previously learned latent space, leading to an *asymmetric* approach that eliminates the need to retrain the NN-based prediction model. Previously proposed asymmetric batch alignment techniques^20^, such as Mutual Nearest Neighbours (MNN)^21^, make stringent assumptions on the nature of batch effects, such as orthogonality between technical variability and biological variability in the data. While these assumptions allow batch alignment through a simple subtraction operation, they do not generalize in the many cases where these assumptions are invalid. In contrast, JIND does not require such assumptions to perform the asymmetric alignment. In addition, JIND estimates cell-type-specific confidence levels during training, which capture the ease to distinguish each type from the rest. These confidence levels are then used to filter out (that is, label as unassigned) cells that cannot be classified with high confidence. Finally, the JIND framework allows the refinement of the parameters of the prediction model via self-training^22, 23^, by treating the high confidence predictions on the target batch as new labeled data. In what follows, we refer to this extension as JIND+.

In summary, JIND is the first automated cell-identification method that provides a scalable framework for accurate label transfer from an annotated source batch to an unannotated target batch, while accounting for existing variability among the two batches. We show that JIND outperforms state-of-the-art methods on most datasets, achieving approximately 97% classification accuracy on average. We also show that the proposed thresholding scheme is robust to datasets of varying difficulties, rejecting only about 4% of cells, while state-of-the-art methods reject considerably higher proportion of cells on average. The misclassification rate can be further reduced with JIND+.

## Results

JIND tackles the problem of supervised cell-type annotation of single-cell RNA sequencing data. The label information comes from a *source batch* dataset: a gene expression matrix with *N_s_* cells (rows) and *M* genes (columns), and the corresponding cell-type annotations (Figure 1(a1)). The goal is to label another dataset, referred to as the *target batch*, which contains the gene expression of *N_t_* cells for the same *M* genes, but no cell-type information. While existing methods require separate batch alignment techniques to be performed prior to classification, JIND trains a NN-based prediction model on the annotated source batch and then uses adversarial training to align the target batch onto the latent space learned by the NN. Thus, JIND is able to compensate for batch effects while avoiding the need for retraining the model when new data becomes available.

**Figure 1:**
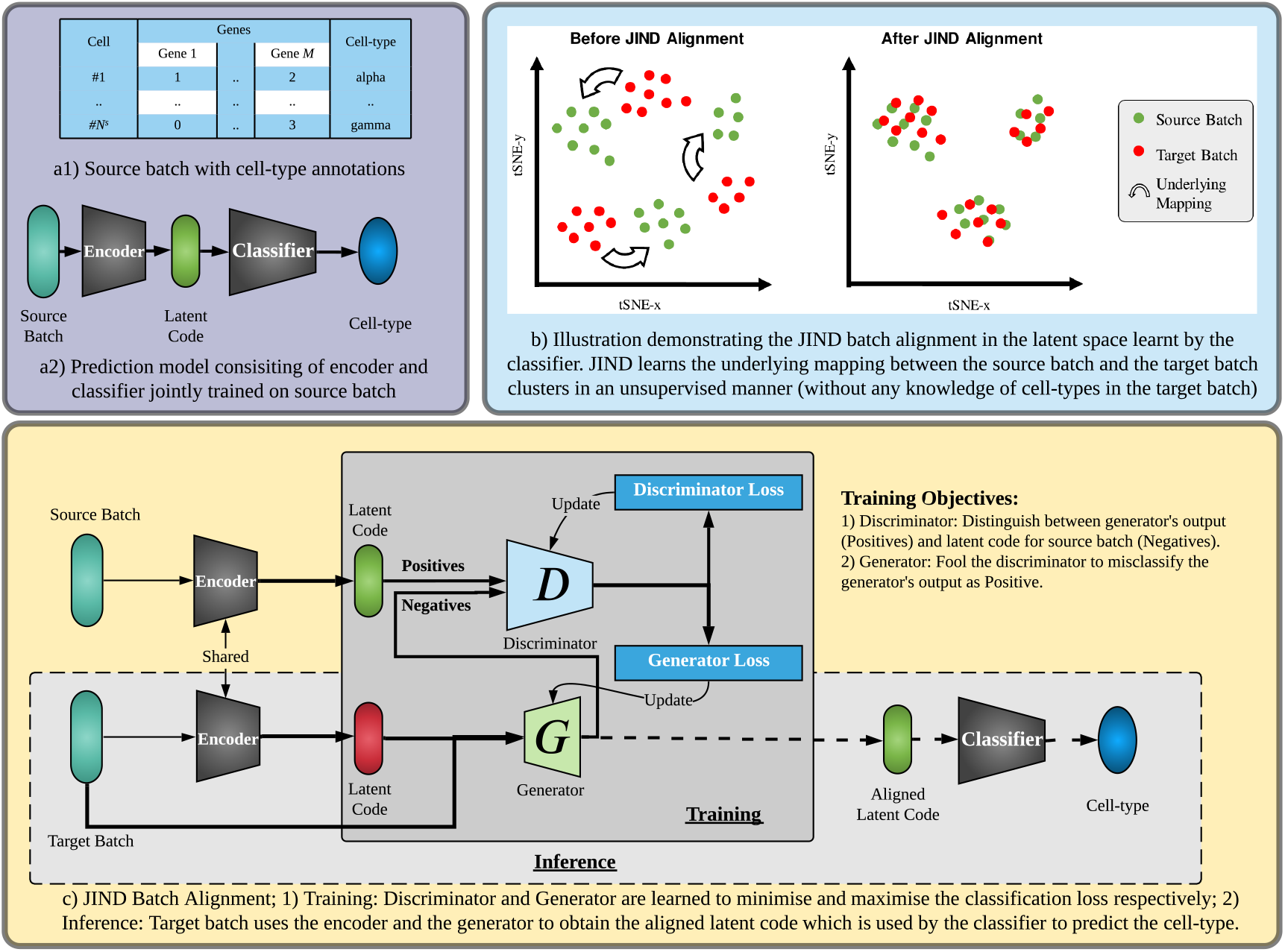
Overview of JIND. a1) We assume access to a source batch containing the gene expression matrix accompanied with the corresponding cell-types. a2) A Neural Network-based prediction model, consisting of an encoder and a classifier, is trained on the source batch. The low-dimensional representation output by the encoder subnetwork is denoted as the *latent code*. Note that this prediction model should not be directly used to annotate the target batch due to batch effects. b) To account for the technical variability across batches, batch alignment is required to align the source and target latent codes. c) JIND uses adversarial training via a generator and discriminator pair to align the source and target latent codes. The discriminator is trained to classify an input latent code either as a latent code produced by the generator (negative label) or as the source latent code produced by the encoder (positive label). In contrast, the generator is trained to fool the discriminator into misclassifying the generator’s output as source latent code. Finally, the output of the trained generator (the aligned latent code) is used by the classifier subnetwork to infer the cell-types of the target batch.

### Overview of the method

The NN used by JIND consists of two subnetworks, an encoder and a classifier. First, the encoder network maps the input gene expression vector onto a 256-dimensional latent space via a one-layer NN. We refer to the resulting 256-dimensional vector as the latent code, which is then fed into the classifier subnetwork to finally predict the cell-type (Figure 1(a2), Supplementary Figure S5). These two subnetworks are trained jointly on the source batch by minimizing a weighted categorical cross entropy loss (see Methods). Since the target batch can have, in general, a different gene expression distribution than the source batch, the latent code (i.e., the encoder output) for both batches will likely have different distributions. Therefore, the latent code from the target batch needs to be modified so that the classifier subnetwork-which was trained on the source batch-can reliably predict the true cell-type (Figure 1(b)).

The proposed alignment technique, which aims at removing batch effects while maintaining useful biological variability for classification, is inspired by both Generative Adversarial Networks (GANs)^24^ and methods developed for the Machine Learning problem known as domain adaptation^25^. More precisely, JIND uses adversarial training to correct the latent code from the target batch by learning a generator function to transform the distribution of the target latent code to that of the source latent code. To learn this generator function, a binary discriminator function is simultaneously learned. While the discriminator function aims at distinguishing between the generator’s output and the source latent code, the generator function aims at fooling the discriminator into misclassifying the generator’s output as source latent code. The output of the trained generator function is the aligned latent code, and it is later used for cell-type inference (Figure 1(c), Methods).

Since it is possible that some cells in the target batch might be undergoing cell differentiation, or that their gene expression might have abnormal patterns, JIND provides a structured way to reject (that is, label them as unassigned) some of the predictions made by the aforementioned prediction model. Specifically, JIND estimates cell-type-specific confidence thresholds from the source batch such that the overall misclassification rate is minimized. This is in comparison to other fixed-threshold-based rejection schemes used in existing methods which do not take into account the variability in ease of classification across different cell-types and datasets (see Methods).

Finally, an extension to the JIND framework based on self-training^22, 23^, coined JIND+, is proposed. In JIND+, additionally, the confident predictions made on the target batch post alignment are used to further fine-tune the parameters of the encoder and classifier subnetworks (see Methods).

### Datasets

In order to assess the performance of the proposed method JIND, we consider five scRNA-seq datasets (Table 1). Specifically, we consider three single-batch datasets: (i) *Human-Hemato*^26^ (GEO Accession ID GSE139369) with 26 annotated cell-types collected from Human blood; (ii) *Mouse Cortex*^27^ (GEO Accession ID GSE60361) with 7 annotated cell-types collected from the mouse brain cortex; and (iii) the *Mouse Cell Atlas*^9^ dataset with 13 annotated cell-types and more than 250 thousand cells.

**Table 1:** scRNA-seq datasets used for evaluation. The *kNN* score indicates the proportion of cells correctly classified using *k*-nearest-neighbour classifier (with *k* = 3), and it serves as an indicator of the dataset complexity (higher kNN score means that cells are easier to classify). For datasets with multiple batches, we report the average *kNN* score across all batches.

| Name [Batches] | Cells × Genes | Cell-types | Batches | Complexity ($kNN$) |
| --- | --- | --- | --- | --- |
| *Human-Hemato*<sup>26</sup> | $35582 \times 20287$ | 26 | × | 0.85 |
| *Mouse Cortex*<sup>27</sup> | $3005 \times 19972$ | 7 | × | 0.97 |
| *Mouse Atlas*<sup>9</sup> | $267690 \times 2797$ | 13 | × | 0.91 |
| *PBMC* [10x_v3, 10x_v5]<sup>28</sup> | $15476 \times 1199$ | 9 | 2 | 0.95 |
| *Pancreas* [Bar16<sup>29</sup>, Mur16<sup>30</sup>, Seg16<sup>31</sup>] | $14058 \times 2448$ | 22 | 3 | 0.88 |

We also consider the following batched datasets: (i) *PBMC* (Peripheral Blood Mononuclear Cells) dataset^28^, containing two batches ‘10x_v3’ and ‘10x_v5’ which differ in the type of assay used during library preparation; and (ii) a *Pancreas* dataset, a collection of three different human pancreas datasets, namely ‘Bar16’ (Baron16^29^), ‘Mur16’ (Muraro16^30^) and ‘Seg16’ (Segerstolpe16^31^). These three pancreas datasets were collected using different sequencing protocols and hence exhibit significant technical variability. As such, they are widely used to benchmark batch correction methods^5^.

We remark that the *Mouse Atlas, PBMC and Pancreas* datasets were already library-normalized and filtered^28^, and only the most informative genes (~3,000) were available at the time of acquisition from ftp://ngs.sanger.ac.uk/production/teichmann/BBKNN. The *Human-Hemato* and *Mouse Cortex* datasets were further processed as described in Methods (Data Preprocessing).

Since most cell-type classification methods have been shown to benefit from external batch correction tools^5^, we integrate the batched datasets using Seurat^15^, obtaining three more datasets, namely, *PBMC Intg, Pancreas Bar-Mur Intg*, and *Pancreas Bar-Seg Intg*.

For every dataset, we also report a *kNN* score that determines the proportion of cells that are correctly classified in the source batch using a *k*-nearest-neighbour classifier (wth *k* = 3) applied to a 20-dimensional representation of the data (obtained via UMAP reduction^32^). This *kNN* score is an indicator of the difficulty/complexity of the dataset and a higher *kNN* score depicts a lower difficulty (Table 1). Note that this score does not take into account the batch effects and therefore does not perfectly correlate with the performance of cell-type identification methods across different datasets (Table 2).

**Table 2:**
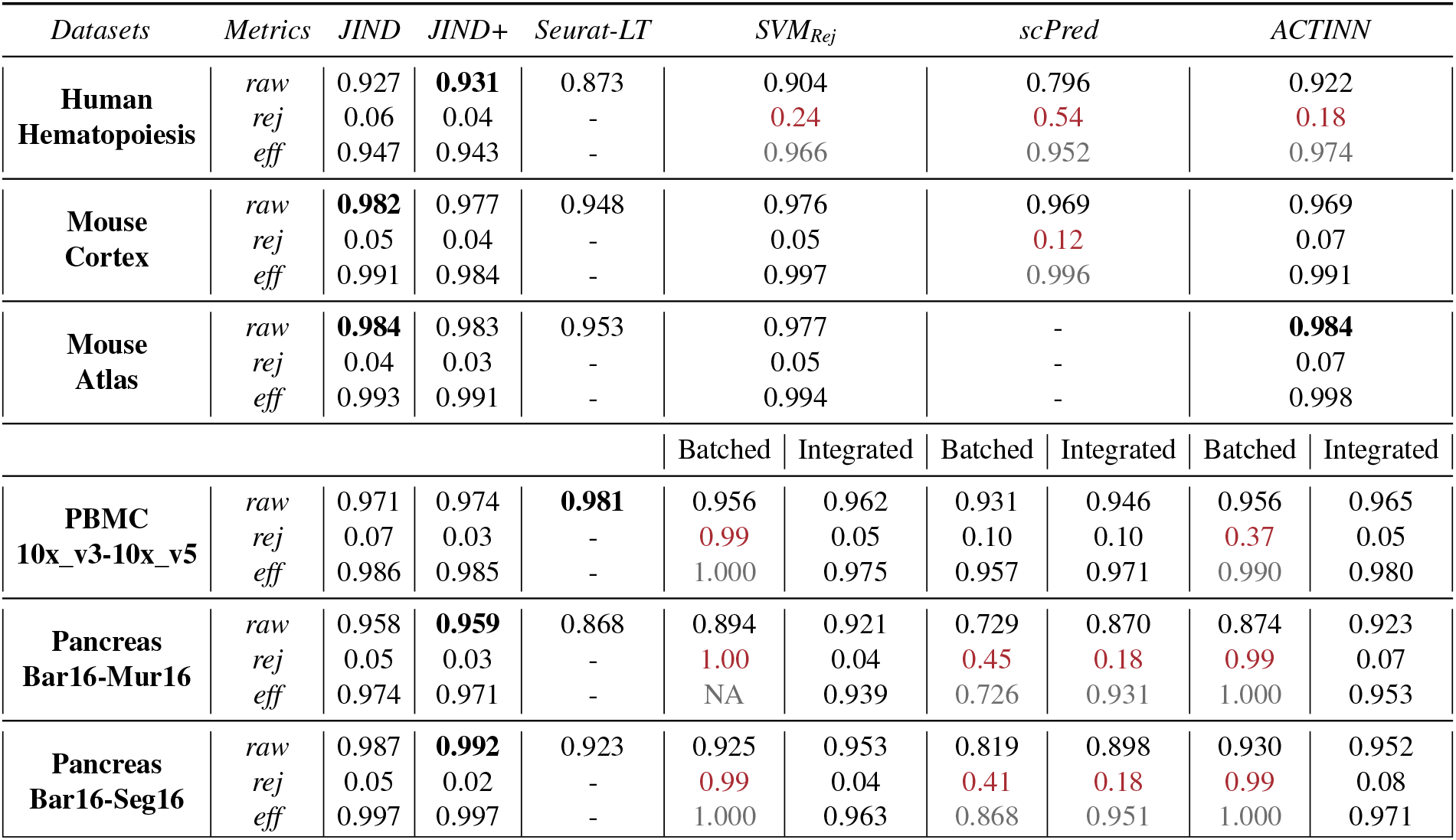
Comparison of different cell classification methods. *raw* is the inital accuracy of the classifier, *rej* is the percentage of cells rejected by the classifier and *eff* is the effective accuracy after rejecting unconfident predictions. For *SVM_Rej_, scPred* and *ACTINN*, we report results without any batch alignment (Batched) and with batch alignment prior to classification using Seurat (Integrated), for the batched datasets *PBMC* and *Pancreas*. Best raw accuracy rates are **bold faced** and rejection rates above 0.1 are colored red.

### Evaluation Metrics

Cell identification methods are commonly benchmarked based on the classification accuracy on the target batch^10, 12, 14^. However, most current methods incorporate an option to reject cells with low-confidence predictions. Thus, it becomes essential to also quantify the proportion of rejected cells, together with the accuracy on the remaining ones. In this context, we use “raw accuracy” (*raw*) to refer to the accuracy of the classifier on the target batch prior to any rejection; “rejection rate” (*rej*) to refer to the percentage of cells rejected by the classification method; and “effective accuracy” (*eff*) to refer to the accuracy of the classifier on the cells that were not rejected by the method.

### Experimental setup

We compare JIND and JIND+ with SVM_Rej_^5^, scPred^11^, Seurat-LT (Seurat Label Transfer)^15^ and ACTINN^10^. These methods were selected as representatives as in a recent study^5^ SVM_Rej_ was shown to perform better than most existing automated cell identification methods as well as methods incorporating prior knowledge in the form of marker genes, and ACTINN^10^ and scPred^11^ were among the best performing methods.

SVM_Rej_ uses a linear Support Vector Machine (SVM) classifier followed by probability calibration using Platt’s method^33^, and sets a confidence threshold of 0.7 to reject cells. scPred also uses a linear SVM classifier but uses only a few PCA (Principal Component Analysis) components with high variance as input features. scPred uses a confidence threshold of 0.9 for rejecting low confidence predictions. Seurat-LT projects the PCA structure (instead of learning a joint CCA structure) of the source batch onto the target batch so as to transfer labels^15^. Seurat-LT does not use any rejection scheme and therefore ‘rej’ and ‘eff’ metrics are not reported for this method. Finally, ACTINN uses a 3 hidden layer NN as the base classifier. We use the confidence threshold of 0.9 for rejecting ACTINN predictions.

In comparison to these methods, JIND is unique in the sense that it learns a low-dimensional representation suitable for cell classification directly from the gene expression data, incorporates a novel asymmetric batch alignment method that projects the unseeen cells into the low-dimensional representation learnt during training, and includes cell-type specific capabilities to detect–and reject–ambiguous cells.

For the datasets that do not contain batch effects, we randomly split the dataset into a 7:3 ratio to generate source and target batches. For the datasets containing batches, we specify the source and the target batch. For example, *Pancreas* Bar16-Mur16 denotes that *Pancreas* Bar16 is the source batch and *Pancreas* Mur16 is the target batch. We note that in *Pancreas* data, there is a wide variation in the set of cell types among the three batches (Bar16, Mur16 and Seg16). Hence for the *Pancreas*-based benchmark experiments on accuracy, and following what was done in the recent review by Abdelaal et al.^5^, while generating the source and target batches we only retained cells whose cell-type is present in both batches.

### JIND achieves low rejection rates with high accuracy on non-batched datasets

While the main benefit of JIND is in the presence of batch effects between the source and target batches, we first perform an assessment on non-batched data. When applied to datasets without a discerning batch effect, JIND and JIND+ achieve a rejection rate of less than 6% for all tested datasets, while maintaining an effective accuracy ranging from 0.945 to 0.99 (Table 2). In addition, they obtain the highest raw accuracy in all cases. On the other hand, previously proposed methods reject a varying proportion of cells. Specifically, on *Human-Hemato* dataset, SVM_Rej_ rejects 24% of cells, scPred 54% and ACTINN 18% (Table 2). In comparison, on the other two datasets, *Mouse Cortex* and *Mouse Atlas*, these methods reject a much lower fraction of cells (between 5% and 12%). We were unable to run scPred on the *Mouse Atlas* dataset (which contains more than 250 thousand cells) and hence no results are reported in this case.

While there is a natural trade-off between rejection rate and effective accuracy on the filtered cells, JIND provides an efficient way of controlling the rejection rate. Specifically, the outlier fraction specified during training (set to 0.05 by default) to estimate the cell-type specific thresholds coincides approximately with the percentage of rejected cells from the target batch, which is on average 5% in all tested datasets. This percentage is further reduced to 3.6% with JIND+. Lastly, Seurat-LT achieves approximately an average raw accuracy that is 4% lower than JIND+ and 3% lower than ACTINN and SVM_Rej_. In comparison to scPred, the performance of Seurat-LT is 8% higher on *Human-Hemtao* dataset and 2% lower on *Mouse Cortex*.

### JIND can accurately annotate scRNA-seq datasets with batch effects

In order to assess the classification performance on batched data, we experiment with three pairs of source and target batches: *PBMC* 10x_v3-10x_v5, *Pancreas* Bar16-Mur16 and *Pancreas* Bar16-Seg16, and compare JIND, JIND+, SVM_Rej_, Seurat-LT, scPred, and ACTINN on these datasets. Since SVM_Rej_, scPred and ACTINN do not internally perform any batch alignment, these methods may benefit from external integration tools^5^. Therefore, we also report their performance after aligning source and target batches using Seurat integration^15^. Table 2 summarizes the results for these experiments. We observe that JIND+ consistently achieves slightly better performance than JIND in all cases. Moreover, JIND+ reduces the rejection rates of JIND by a factor of 2 while keeping the effective accuracy almost identical. We also observe that JIND and JIND+ outperform previously proposed methods in raw accuracy in all cases, except for the *PBMC* dataset in which Seurat-LT achieves a raw accuracy 0.8% higher than JIND+. Nonetheless, in the *Pancreas* datasets, JIND+ outperforms Seurat-LT by 9% on average. When no external alignment is performed the rejection rates with SVM_Rej_, scPred and ACTINN, for all three datasets, are significantly higher than with JIND+. Notably, SVM_Rej_ and ACTINN reject almost all cells in some cases. It should be noted however that scPred is supposed to be used in conjunction with a batch alignment tool when used on datasets with batch effects^11^. When SVM_Rej_, scPred and ACTINN were evaluated after using Seurat batch alignment, we observe that their rejection rates are significantly reduced. However, scPred still rejects more than 10% of cells even after batch alignment in all three experiments. Our results are in agreement with the review conducted by Abdelaal et al.^5^, which concludes that SVM_Rej_, scPred and ACTINN benefit from batch alignment tools. However, JIND+ still outperforms ACTINN and SVM_Rej_, achieving approximately 3% higher raw accuracy. In comparison to scPred, JIND+ achieves more than 7% percent higher raw accuracy on average.

To further demonstrate the benefits of JIND cell identification and visualize the asymmetric batch alignment, we consider the *Pancreas* Bar16-Mur16 dataset, where *Pancreas* Bar16 is the source batch and *Pancreas* Mur16 the target batch, and retain only the cells with cell-types *Alpha, Beta, Gamma* and *Delta*. The considered source and target batches exhibit profound batch effects, and their latent codes on the tSNE space after JIND’s prediction model is trained on the source batch appear completely segregated (Figure 2(a)). As a result, when used without any batch alignment, JIND’s prediction model achieves an effective accuracy close to 98.7%, but rejects approximately 50% of the cells due to low confidence in the predictions (Figure 2(b)). Running JIND’s asymmetric alignment results in a partial mapping of the latent space between the distinct clusters of the target batch and those of the source batch in the tSNE space (Figure 2(c)). Despite not observing a perfect overlap between the two batches, JIND still rejects less than 5% of the cells and obtains high accuracy on all four cell-types (Figure 2(d)).

**Figure 2:**
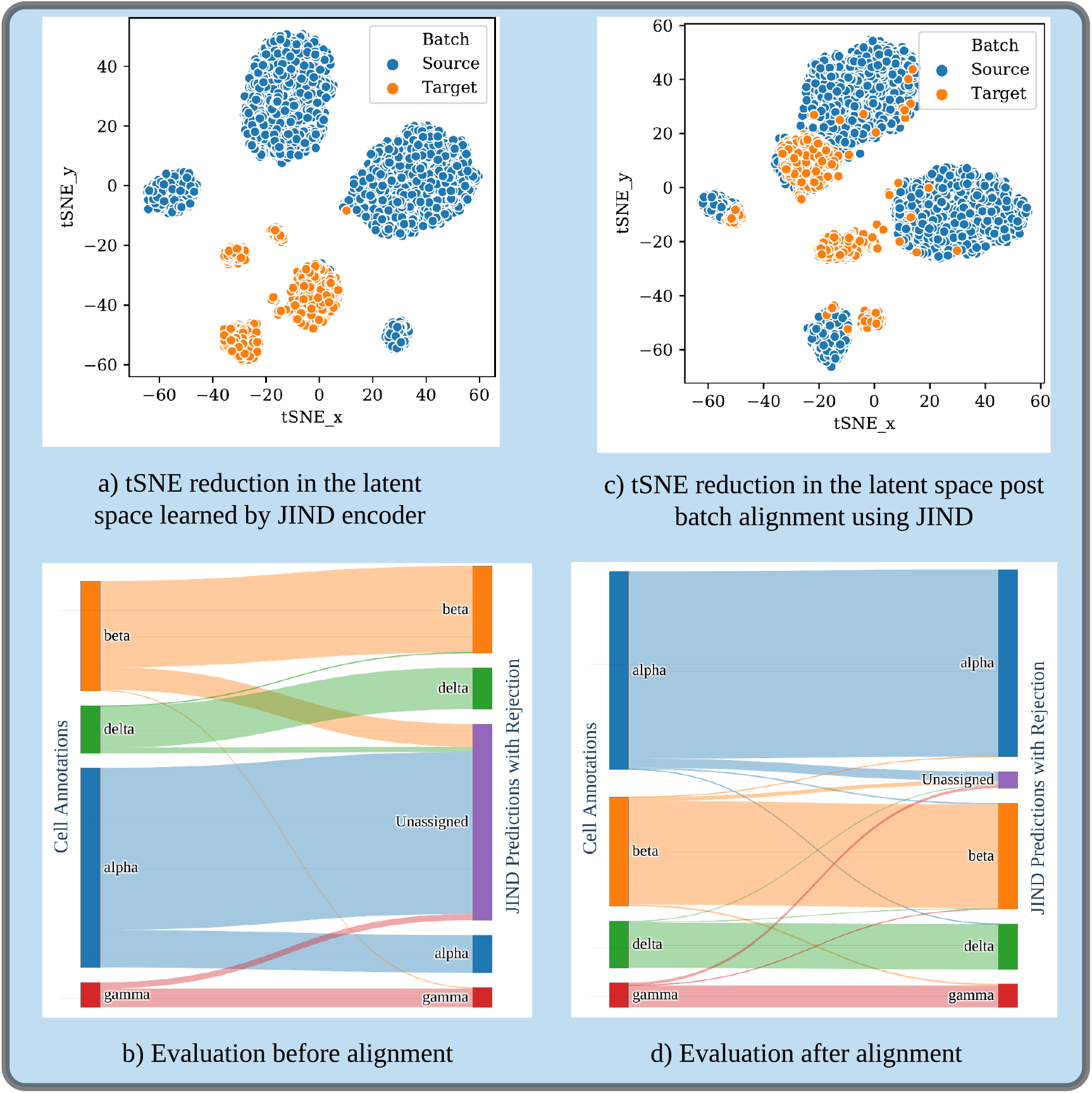
JIND’s asymmetric alignment leads to accurate annotations on batched data. We consider a subset of cell-types (*Alpha, Beta, Gamma* and *Delta*) from *Pancreas* Bar16 (source batch) and Mur16 (target batch). a) tSNE reduction in the latent space shows significant distributional mismatch due to batch effects. b) As a result, the alluvial plot shows that the prediction model (without alignment) makes a large number of “unassigned” predictions. c) JIND batch alignment removes these batch effects using adversarial training (learning the Generator and Discriminator parameters), which minimizes the distributional discrepancies among the two batches in the latent space learned by the encoder subnetwork. d) The alluvial plot thus obtained after performing batch alignment on target batch shows accurate classification performance per cell-type.

While most batch correction methods^15, 20, 34^ aim at achieving perfect overlap between source and target batches, our results demonstrate that JIND does not necessarily require overlap to accomplish the task of classification (Figure 2(c) and Figure 2(d)). For completeness, we also ran JIND and JIND+ after Seurat integration and observed that using Seurat integration actually deteriorates classification performance (Supplementary Table S1). This further shows that JIND is more effective than all existing methods for cell-type identification.

#### Runtime comparison

Since the JIND framework is based on NNs that are inherently parallelizable, JIND can exploit multiple CPU cores for faster runtime and can also be run on GPUs. We compared the runtime for Seurat integration and JIND asymmetric alignment on *PBMC* 10cx_v3-10x_v5 experiment. JIND was able to complete the batch alignment roughly 7 times faster than Seurat on 40 CPU cores (Intel(R) Xeon(R) CPU E5-2698 v4 @ 2.20GHz). Seurat was parallelized using the *future* package (http://CRAN.R-project.org/package=future) and was run on 40 cores as well. Note that, besides faster alignment, JIND framework (being asymmetric) can also amortize prediction model training time in the scRNA-seq dataset annotation pipeline across multiple batches. We also compared the running time of JIND+ including classifier training, batch alignment and self training with scPred, ACTINN, SVM_Rej_ and Seurat-LT under same hardware settings on the *PBMC* 10x_v3-10x_v5 experiment. We observed that both SVM_Rej_ and ACTINN are roughly 2-3 times faster than JIND+ since there is no batch alignment or self-training phase. Seurat-LT was observed to be 75% slower than JIND+ and it includes a batch alignment phase. Lastly, scPred (on one CPU core) was observed to be approximately 3-4 times slower than JIND+. The implementation provided by the authors does not support multithreading. Therefore, JIND is a fast and more accurate alternative for existing automated cell annotation tools.

Since JIND+ either outperforms JIND or achieves similar performance, in the remainder of the paper we will only discuss the performance and results for JIND+.

### JIND+ rejection filters ambiguous cells and improves annotation accuracy

When automatically annotating a target batch using a previously trained classification model, the classification of a cell from the target batch may be ambiguous. Classification ambiguity happens mostly when either the cell lies in a transitioning state, which is usually depicted as the intersection of two clusters of cells in the tSNE-reduced space; or it lies at the boundary of its corresponding cluster (i.e., far from the cluster centroid) in the tSNE-reduced space. Since cells in either of these cases are prone to misclassification, it is important to reject (or flag) them during the annotation process, as manual annotation of such cells may be preferred. Otherwise, wrong annotation of cells might hamper the downstream analysis of scRNA-seq data.

In order to assess JIND+ rejection performance, we consider two datasets, namely *Pancreas* Bar16-Mur16 and *PBMC* 10x_v3-10x_v5. In the case of *Pancreas* Bar16-Mur16 dataset, we observe that most of the unassigned cells are at the intersection of either one of the *Alpha, Delta* and *Gamma* clusters in the tSNE reduced space (Figure 3(a)). Moreover, there exists some *Acinar* and *Delta* cells on the left side of the *Alpha* cluster which were at first misclassified (prior to the rejection step) but later assigned an “unassigned” label. We also observe some outliers such as *Delta* cells in the *Beta* cluster or *Ductal* cells in the *Acinar* cluster, which were originally correctly classified but were later rejected and appear to be outliers in the tSNE space (due to potential mislabeling). In summary, out of 61 cells that were rejected (i.e., labeled as “unassigned”), 26 of those cells were originally misclassified raising the 95.9% raw accuracy to an effective accuracy of 97.1%.

**Figure 3:**
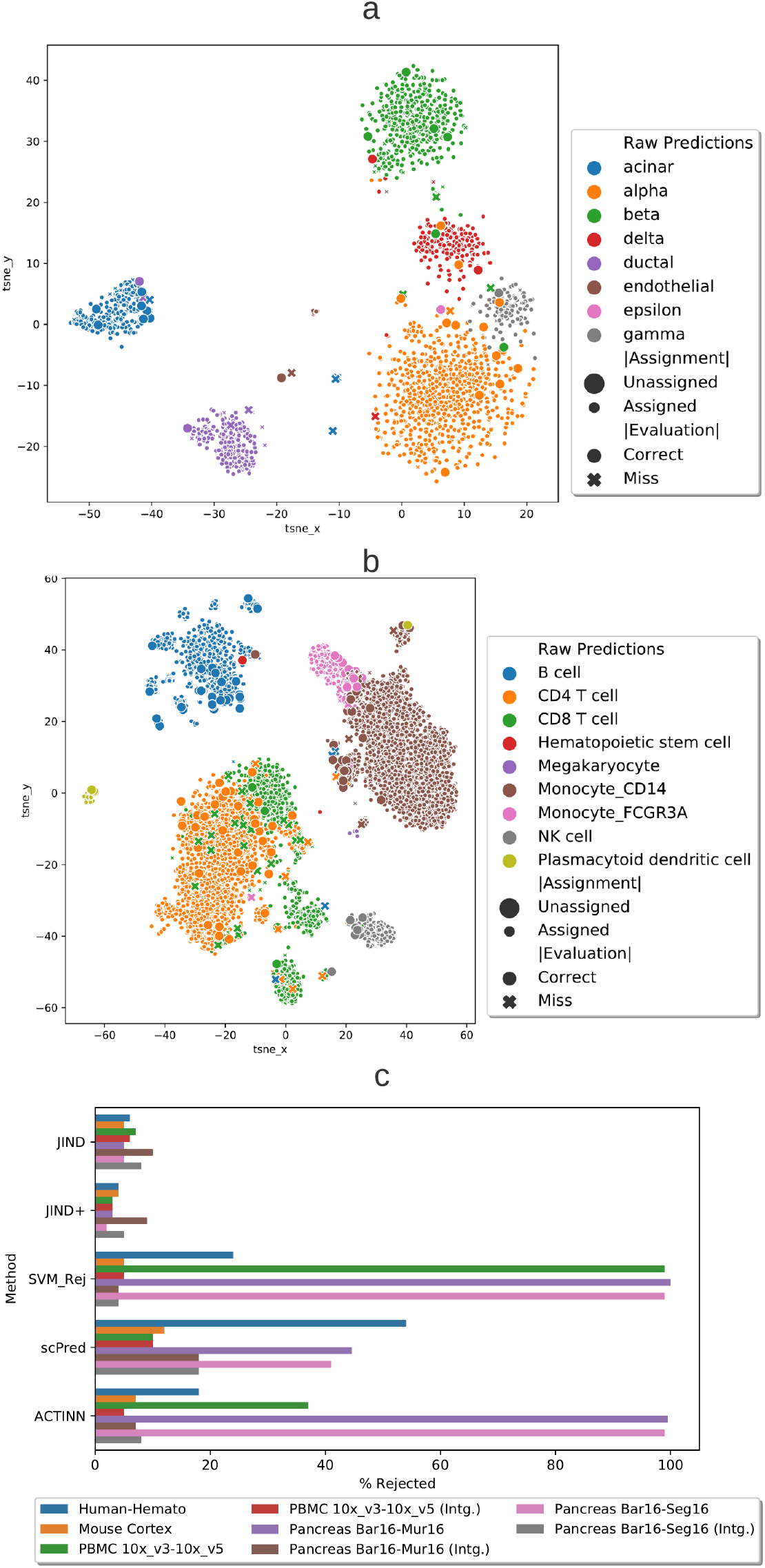
Visualization of cells rejected by JIND+. Cells with their corresponding raw predictions (specified by different colors) in the tSNE space are shown for a) *Pancreas* Bar16-Mur16 and b) *PBMC* 10x_v3-10x_v5 datasets. Incorrect predictions are denoted with a cross, and correctly ones with a filled circle. Many of the rejected cells (specified by a bigger marker size) lay at the intersection of two or more clusters or at the boundary of their corresponding cluster, far away from the centroid. This suggest that they are either transitioning cells, noisy cells (outliers) or mislabeled cells. c) Comparison of the percentage of cells rejected by each method over all considered datasets. We also include in the comparison the batched datasets after integration with Seural (denoted by *Intg*.).

When using the *PBMC* 10x_v3-10x_v5, we observe that a large number of rejected cells lie at the intersection of *Monocyte FCGR3A* and *Monocyte CD14* clusters or the *CD4* and *CD8 T* cell clusters. We also observe that many *CD8 T* cells that lie inside the *CD4* cluster were rejected and initially misclassified by JIND+. This set of *CD8 T* cells lying inside the *CD4* cluster explains why many *CD4 T* cells inside the *CD4* cluster are rejected, as some *CD8 T* cells exist on the left of the *CD4* cluster. Moreover, many of the *B* cells lying on the boundary of the *B* cluster were rejected including an outlier *Hematopoietic* stem cell and an outlier *Monocyte_CD14* cell (Figure 3(b)). In summary, out of a total of 240 cells that are rejected by JIND+, 85 are misclassifications which raises the 97.4% raw accuracy to an effective accuracy of 98.5%.

Finally, we analyze the percentage of rejected cells by each of the considered methods on the datasets without batch effects (*Human-Hemato, Mouse Cortex*) and the datasets with batch effects (*PBMC* 10x_v3-10x_v5, *Pancreas* Bar16-Mur16, *Pancreas* Bar16-Seg16), the latter ones with and without batch integration (Figure 3(c)). As already mentioned, JIND and JIND+ reject a nearly constant small percentage of cells across all datasets (below 10%), with JIND+ having lower rejection rates (about 5%). On the contrary, we observe that on datasets with batch effects, scPred, SVM_Rej_ and ACTINN reject a large fraction of cells, ranging from 10% to 100%. On datasets without batch effects and on the Seurat-aligned datasets, SVM_Rej_ and ACTINN reject less than 10% of cells except on *Human-Hemato* dataset. Since the number of cell-types in *Human-Hemato* is 26, this result is not surprising, as the predicted probability vectors are expected to have high entropy, which translates into the highest probability being below the fixed threshold of 0.9. Finally, scPred rejects a large proportion of cells on all datasets except on *PBMC* 10x_v3-10x_v5 (integrated using Seurat) where it rejects 10% of cells. In conclusion, the rejection mechanism of JIND is superior to the fixed threshold used by previously proposed methods.

### JIND+ misclassified cells exhibit differentially expressed genes

To better understand the misclassifications made by JIND+, we further analyze the results obtained from *Pancreas* Bar16-Mur16 and *PBMC* 10x_v3-10x_v5 datasets. When using the *PBMC* 10x_v3-10x_v5 dataset, we observe that JIND+ misclassifies approximately 1.5% cells after rejection (Figure 4(a)-top). To identify which cell-types can result in misclassifications due to cluster overlaps, we also visualize the target batch (*PBMC* 10x_v5) using tSNE dimensionality reduction (Figure 4(a)-middle). We observe that two subpopulations of *Monocytes*, namely, *Monocyte FCGR3A* and *Monocyte CD14*, lie close to each other with a noticeable overlap in the tSNE-reduced space. Since some of the *Monocyte FCGR3A* cells are misclassified by JIND+ as *Monocyte CD14*, we conduct a differential expression (DE) analysis using Limma^35^ between the misclassified (*Monocyte FCGR3A* predicted as *Monocyte CD14*) and the correctly classified (*Monocyte FCGR3A* predicted as *Monocyte FCGR3A*) *Monocyte FCGR3A* cells.

**Figure 4:**
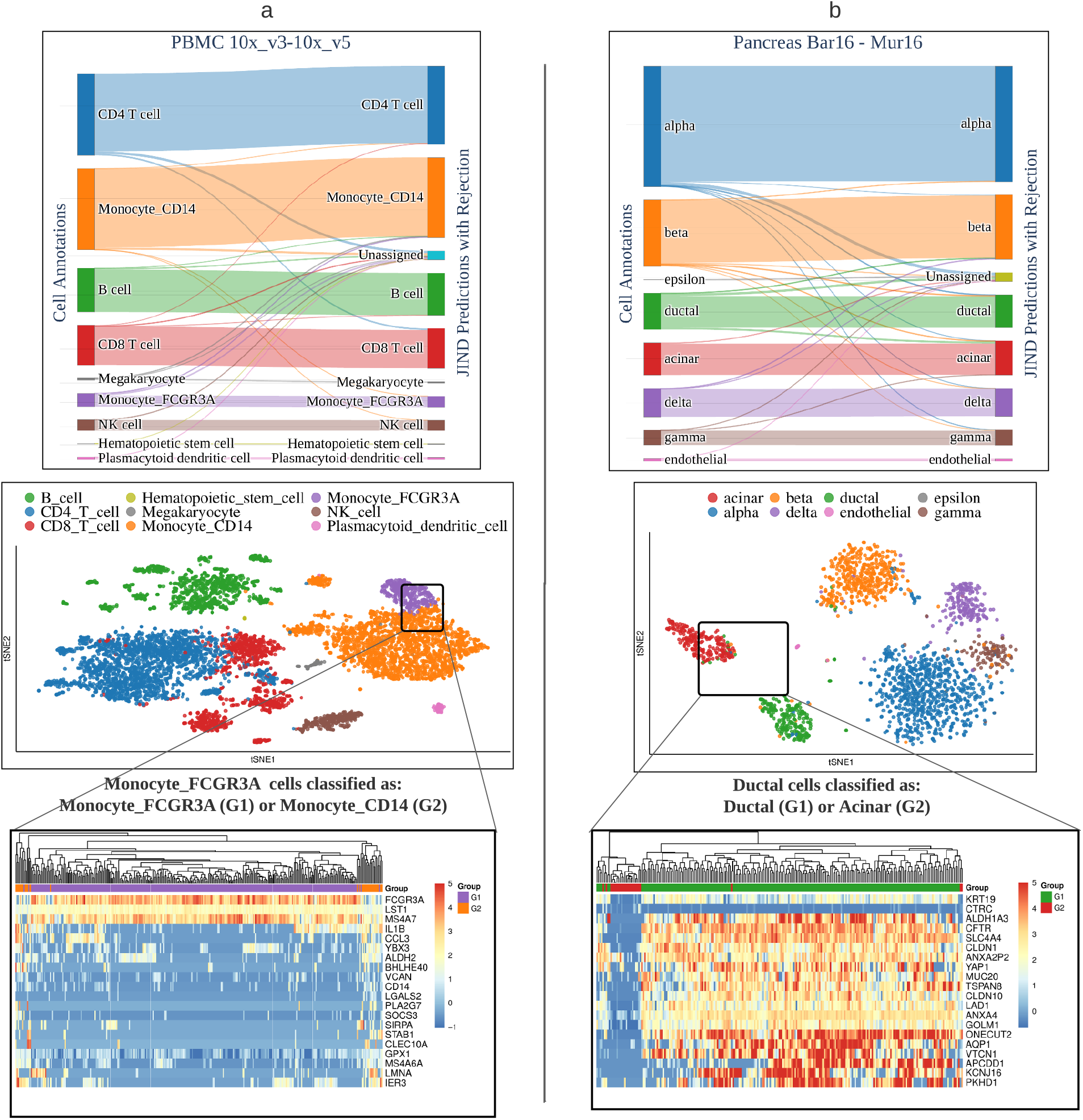
Performance evaluation and differential expression analysis on two datasets. The alluvial plots (top) reflect the performance of JIND+ on a) *PBMC* 10x_v3-10x_v5 and b) *Pancreas* Bar16-Mur16 datasets. The tSNE plots (middle) illustrate the cell-type clusters of the target batch, and highlight the two cell-types with the highest misclassification rates: a) *Monocyte_FCGR3A* and *Monocyte_CD14* and b) *Acinar* and *Ductal*. The heatmaps (bottom) show the top 20 differentially expressed genes between a) *Monocyte_FCGR3A* cells classified as *Monocyte_FCGR3A* (G1) and *Monocyte_FCGR3A* classified as *Monocyte_CD14* (G2), and between b) *Ductal* cells classified as *Ductal* (G1) and *Ductal* cells classified as *Acinar* (G2). The shown hierarchical clustering is performed using all the differentially expressed genes.

We identify 117 significantly differentially expressed (*p* < 0.001) genes between correctly predicted Monocyte *FCGR3A* cells and Monocyte *FCGR3A* cells predicted as *CD14* (Figure 4(a)-bottom, Figure S2, Supplementary Excel File). Furthermore, we observe that the gene marker *FCGR3A* is clearly overexpressed on the group of cells classified by JIND as *Monocyte FCGR3A*. Similarly, we can observe the overexpression of the *CD14* gene on the cells classified as *Monocytes_CD14* and an underexpression on the ones classified as *Monocytes_FCGR3A*. This suggests that the cells misclassified by JIND+ are actually outliers (or possibly mislabelled in the original dataset), thus being intrinsically difficult to classify. Therefore, these misclassifications do not correspond to arbitrary mistakes made by the prediction model.

We perform a similar analysis on the *Pancreas* Bar16-Mur16 dataset. We observe that JIND+ misclassifies roughly 2.9% of cells from the *Pancreas* Mur16 dataset after rejection (Figure 4(b)-top). We again visualize the target batch using tSNE dimensionality reduction to identify cells that are hard to classify. Interestingly, JIND+ misclassifies about 5% of the *Ductal* cells as *Acinar*, even though the clusters do not overlap in the tSNE space (Figure 4(b)-middle). On close observation, we find that some of the *Ductal* cells actually lay closer to the *Acinar* cluster centroid than the *Ductal* centroid. Hence, we perform a DE analysis between the misclassified (*Ductal* predicted as *Acinar*) and correctly classified *Ductal* cells (*Ductal* predicted as *Ductal*). The analysis reveals that 444 genes are differentially expressed (*p* < 0.001), among which we also find biomarkers for the two cell-types (Figure 4(b)-bottom, Figure S1, Supplementary Excel File). Specifically, *KRT19*, a positive biomarker gene^36^ for *Ductal* cell-type, is significantly underexpressed on the misclassified group and *CTRC*, a positive biomarker for *Acinar* cells^36^, is differentially expressed but with a very low expression pattern. These findings suggest that the *Ductal* cells classified as *Acinar* are indeed not *Ductal*, and that further analysis is needed to confirm their true identity.

We observe that the number of misclassifications is less than 50 for both experiments in total. Therefore, we also conduct a DE analysis by randomly selecting a subset of cells (of the same size) from *Monocyte FCGR3A* cells as well as *Ductal* cells. We observe that any two random subsets of cells do not exhibit differentially expressed genes and therefore cannot be distinguished based on gene expression profiles (Supplementary Figure S3 and S4, Supplementary Excel File). We conclude that some misclassifications made by JIND+ are explainable and likely due to issues with the data labels.

### JIND batch alignment learns a meaningful mapping in extreme cases

One of the inherent limitations in transferring cell-type information from a source batch to a target batch under significant batch effects is that batch alignment or integration becomes extremely hard when the cell composition of the two batches is different. Specifically, if there exist cell-types in the target batch that are not present in the source batch, then alignment in general becomes an ill-defined problem. Moreover, without a priori knowledge, the new cell-type in the target batch would likely be misclassified, as the alignment method in such a scenario might result in a false positive matching of cell-types^34^.

To investigate this point, we analyze how JIND asymmetric alignment (in the latent space) maps the clusters in a controlled setting with new cell types in the target batch. We consider the *Pancreas* Bar16-Mur16 dataset and select from Bar16 the *Alpha, Beta, Gamma* and *Delta* cell-types to generate the source batch, and from Mur16 the *Acinar* cell-type along with the four cell-types in the source batch to generate the target batch. As expected, after training JIND’s prediction model on the source batch, the latent codes for the target and source batches in the tSNE space reveal four and five clusters in the source and target batches, respectively (Figure 5(a)). Moreover, due to batch effects, no overlap between any of the clusters is observed. We then run JIND’s asymmetric alignment and infer cell-types for the target batch. Interestingly, while many *Acinar* cells are rejected, most of them are classified as *Beta* (Figure 5(b)). Visualizing the aligned latent codes for the source and target batches using tSNE dimensionality reduction shows that three target batch clusters are clearly mapped to three source batch clusters (Figure 5(c)). We enumerate the remaining two clusters from the target batch as Cluster 1 and Cluster 2. In the source batch, only the *Beta* cluster is not mapped to any one cluster of the target batch. Therefore, we compute the average Euclidean distances in the latent space from Cluster 1 and Cluster 2 to the *Beta* cluster in the source batch. We observe that *Beta* cells from the source batch are actually twice as close in the latent space to Cluster 2 than to Cluster 1, implying higher biological similarity between Cluster 2 and *Beta* cells. Cross-checking with cell annotations, we find that Cluster 1 corresponds to *Acinar* cells and Cluster 2 to *Beta* cells (Figure 5(d)). This shows that even in cases with new cell-types in the target batch, the mapping learned by JIND’s asymmetric alignment is meaningful and captures biological similarity between cells. It must be noted however that it is very hard in general to detect novel cell-(sub)types present in the target batch, as in most cases the clusters are not well separated in the latent space, making such analysis much more difficult.

**Figure 5:**
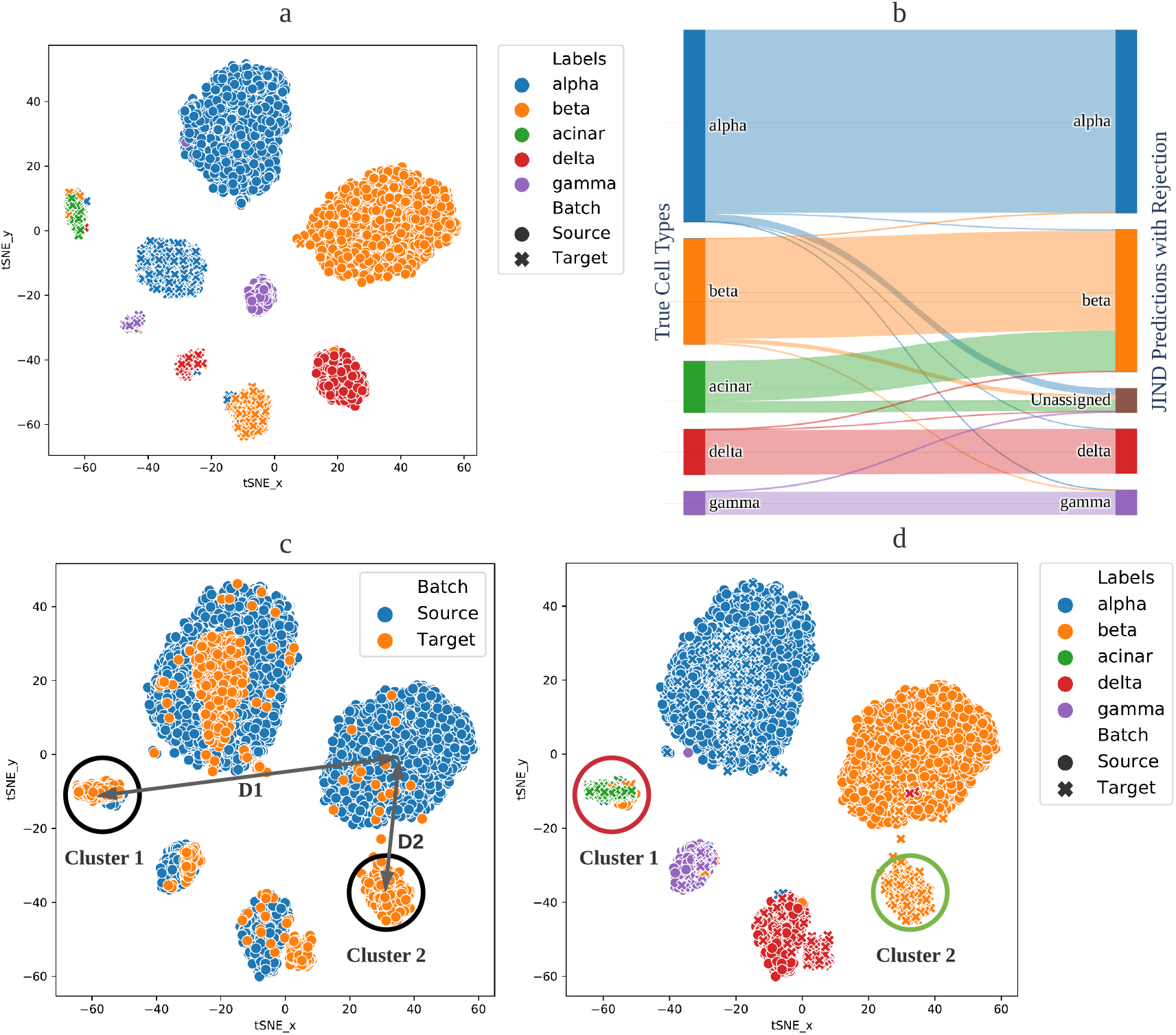
Analysis of the JIND asymmetric integration when a novel cell-type is present in the target batch. We consider the *Pancreas* Bar16-Mur16 dataset, but select only the cells in the source and target batches that are annotated as *Alpha, Beta, Gamma* or *Delta* and additionally the *Acinar* cells in the target batch. a) Before alignment, the four cell-types from the source batch and the five cell-types from the target batch are shows as isolated clusters in the tSNE-reduced space. b) After JIND’s asymmetric alignment and classification, we observe that most of the novel *Acinar* cells are labeled as *Beta*, but a significant fraction of them is labeled as “unassigned”. c) After JIND’s asymmetric alignment, three of the clusters in the target batch are properly aligned in the tSNE space to the source batch clusters, while the remaining two are not. By computing the distance between these two target batch clusters (denoted as Cluster 1 and Cluster 2) and the unmapped cluster in the source batch, we notice that Cluster 2 is much closer to the unmapped cluster. d) We observe that the unmapped source cluster corresponds to *Beta* cells, and that the target Cluster 1, which is further from the *Beta* cluster is indeed the novel cell-type *Acinar* introduced in the target batch. This suggests that a careful distance analysis in the JIND latent space can help disambiguate the mapping between source and target batches in the presence of new cell-types.

## Discussion

In this work we introduced JIND, an automated cell-type identification tool that utilizes pre-annotated scRNA-seq data to reliably annotate unseen sequenced data. Depending on the differences in sequencing protocols or data preparation, there typically exists significant technical variability between these datasets, which confounds real biological variability. To accurately predict cell-types while dealing with potential batch effects, JIND uses supervised learning in addition to adversarial training to train a prediction model as well as learn a mapping to align these datasets. The prediction model and the mapping are then used in conjunction to infer cell-types of unseen data. The mapping is learned after training the prediction model and hence any unseen dataset can be annotated directly without the need of retraining the prediction model. This is in contrast to other cell-type identification methods that rely on symmetric batch integration tools and require training the prediction model after performing batch alignment. In addition, the prediction model used by JIND is based on neural networks (NN) and directly learns the most informative features for cell classification from the scRNA-seq data, eliminating the need of prior feature extraction. JIND also incorporates a robust rejection scheme which filters out low-confidence predictions to avoid the misclassification of cells in ambiguous states or that have highly noisy gene expressions. This is done by estimating cell-type-specific confidence thresholds which are determined using the annotated data, and therefore adapt to the dataset complexity. Lastly, we presented JIND+, and extension of JIND that uses the confident predictions made on the unseen data to further fine-tune the parameters of the prediction model.

We demonstrate that both JIND and JIND+ achieve higher accuracy on cell-type identification for datasets containing batch effects as compared to existing state-of-the-art methods. This is accomplished while maintaining a constant rejection rate (about 5%) which can be easily controlled by the user. We investigated the misclassifications made by JIND+ on two datasets and observed that they can be explained by variabilities on the expressions of the cell-type biomarkers (genes). We also showed that the cells rejected by JIND generally correspond to misclassified cells, improving the effective classification accuracy by reducing the misclassification rate. In conclusion, the observed improvements in the performance on cross-batch annotation demonstrate that JIND is highly effective at aligning batches and discriminating cell-types.

## Methods

JIND is a framework for automatic cell identification based on supervised learning. Given a scRNA-seq dataset with annotated cells (denoted as the source batch), the goal is to train a model that can then be used to predict the cell annotations of an independently generated scRNA-seq dataset for which cell annotations are not available (denoted as the target batch). We assume that both datasets contain roughly the same set of cell-types.

Next we outline the different components of JIND, and describe JIND+, an extension of JIND that additionally employs self-training to fine-tune the model parameters.

### Data preprocessing

scRNA-seq data can be expressed as a matrix **X** of dimension *N* × *M*, with *N* and *M* denoting the number of cells and genes, respectively. We denote the source batch used for training the prediction model by **X**^s^ (*N_s_* × *M*), and the corresponding cell annotations by **Y**^s^. We assume the cell annotations are represented as hot-encoded *K*-dimensional vectors, where *K* is the number of cell-types in the source batch, such that **Y**^s^ ∈ {0, 1}^*N_s_*×*K*^. For example, if *K* = 5 and a given cell belongs to the third class, the corresponding row of **Y**^s^ is encoded as (0, 0,1, 0,0). At the prediction step, we denote the gene expression matrix for the target batch by **X**^t^ (*N_t_* × *M*). No cell annotations are available for the target batch.

The neural-network-based prediction model implicitly learns the appropriate representations^37^ relevant for performing the classification task directly from **X**^s^. Nevertheless, since scRNA-seq data is high-dimensional (it may contain the expression of approximately 20k-40k genes^38^) and most counts are near zero, some preprocessing is needed. In particular, we first apply the standard log-transform to the source expression matrix using log_2_(1 + **X**^s^), as done in related works^10, 11^. Then, we select the top 5000 genes exhibiting the highest cell to cell variation, similar to what is done by Seurat^15^. In the case when fewer than 5000 genes are available, all genes are selected for training. JIND selects 5000 genes by default, as it provides a good trade-off between complexity and classification accuracy (Supplementary Table S2). Note that this hyperparameter can be easily modified in the JIND framework. The log-transformed values of the selected genes are used to train the prediction model, together with the cells’ annotation information **Y**^s^. With some abuse of notation, in the rest of this manuscript, we refer to the log-transformed values of the selected genes as **X**^s^. Similarly, during inference, **X**^t^ will denote the log-transformed expressions of the same subset of genes that were selected during training.

### Training stage

#### Prediction model

The prediction model used in JIND is based on NNs and consists of two subnetworks (Supplementary Figure S5): (i) an *encoder*, which contains one hidden layer with 256 neurons, and (ii) a *classifier* consisting of one hidden layer with 256 neurons followed by a softmax layer which outputs K probabilities (with K being the number of distinct cell-types in the source batch). Both subnetworks employ ReLU^39^ (Rectified Linear Unit) non-linearity as the activation function, which is typically used in deep NNs. The output of the encoder prior to the ReLU activation function is referred to as the *latent code*. The hidden layer in the encoder subnetwork uses dropout^40^ to avoid over-fitting while training, with a dropout probability of 0.2, as in other cell-type identification methods^10, 41^. The output of the prediction model is a *K*-dimensional vector *ŷ* representing the probabilities of the cell belonging to each of the cell-types.

The network parameters are trained by minimizing the weighted categorical cross-entropy loss. We denote the expression data for one cell by the vector *x* containing the expression for *M* genes (5000 by deafult); and the corresponding cell annotation, encoded as a one-hot encoded K-dimensional vector, by *y*. For input *x*, the weighted categorical cross-entropy loss is defined as

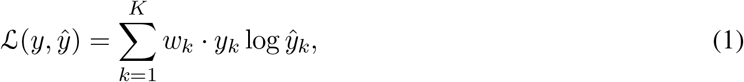

where *y_k_* and *ŷ_k_* denote the *k*th entry of the vectors *y* and *ŷ*, respectively, and *w_k_* is a constant scalar determined as a function of the proportion of cells annotated as the *k*th type. A weighted loss is used to account for a potential class imbalance in the dataset. The weights in the objective function are inversely proportional to the number of cells in the training dataset belonging to that cell-type. If *n_k_* denotes the fraction of the *k*th cell-type in the dataset, then the weights can be calculated as

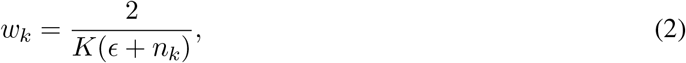

where *ϵ* is added to the denominator to avoid assignment of an exceedingly high weight to a cell-type, and is set to 0.01 by default.

#### Training Details

We split the source dataset (**X**^s^, **Y**^s^) into training (70%) and validation sets, and then train the NN model for 30 epochs on the training dataset. After each epoch, the resulting model is evaluated on the validation set, and the corresponding validation accuracy is recorded. At the end of the training process, the model parameters corresponding to the highest validation accuracy (i.e., the smallest misclassification rate) across all epochs are selected and saved. Although the training loss is expected to decrease as more epochs are completed, the validation accuracy may decrease, for example if over-fitting occurs. An Adam optimizer^42^ with an initial learning rate of 0.001 is used to optimize the network parameters. The learning rate is reduced by a factor of two if the training loss saturates for more than 5 epochs.

#### Filtering

To make the misclassification rate minimal, JIND uses *K* confidence thresholds, one per cell-type, denoted by *τ_k_*, with *k* ∈ [1: *K*]. At inference time, before mapping a cell to the *k*th cell-type corresponding to the highest probability, we cross check whether the probability is greater than the corresponding threshold *τ_k_*, resulting in an “unassigned” label upon failure. The thresholds are determined as a part of the training process. We use the validation dataset to select a threshold for each cell-type based on an outlier fraction *θ* (set to 0.05 by default). Specifically, the threshold *τ_k_* is the highest predicted probability of the bottom *θ*-quantile of the cells assigned to *k*th cell-type.

### Inference stage

Once the training of the prediction model has been completed, the next step is to use it to classify the cells of an independent scRNA-seq dataset, referred to as the target batch **X^t^**, for which the cell annotations are not available. In most practical scenarios, however, the target batch may exhibit batch effects, which translate into differences between the distributions of **X**^s^ and **X^t^**. To account for these potential differences and to avoid misclassification, JIND uses a novel and scalable asymmetric alignment technique based on adversarial training that aligns the target batch onto the source batch.

#### Asymmetric integration

JIND aligns the target batch onto the source batch in the latent space learned by the encoder subnetwork after training the NN-based prediction model (Supplementary Figure S5). This alignment is attained by transforming the latent code obtained from the encoder subnetwork to the target batch so that it is indistinguishable from the latent code obtained from the source batch.

Let the function learned by the encoder subnetwork (prior to applying ReLU activation) be represented by *F* and the one learned by the classifier subnetwork by *P* (which includes the ReLU activations after the encoder subnetwork). Thus, the predictions are produced as *P*(*F*(*x*)), where *h* = *F*(*x*) are the corresponding hidden representations obtained from *F* (the latent code). Our task is to learn a function *G*(*x, h*) such that *h* = *F*(*x*) with *x* ~ **X**^s^ (source batch), is indistinguishable from *ĥ* = *G*(*x*, *F*(*x*)) with *x* ~ **X**^t^ (target batch). Once such function *G* is found, we expect *P*(*ĥ*) to produce accurate cell-type predictions. We perform the asymmetric integration in the latent code prior to ReLU activation as we observed that this choice led to improved performance (Supplementary Table S3). We assume the following functional form for the function *G*, referred to as the generator, that shifts and scales *h* to obtain the modified latent code *ĥ*:

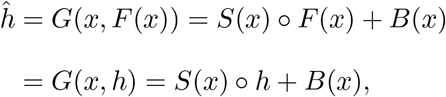

where ∘ indicates element-wise multiplication, and *S* and *B* are two different NNs jointly parameterized by Θ_*G*_ that scale and shift *h*, respectively. This mapping is motivated by residual connections^43^, which have been shown to be extremely successful at learning deep NNs. Since the latent code *h* produced by the encoder subnetwork is a 256-dimensional vector, the outputs of the NNs *S* and *B* are also of dimension 256.

#### Adversarial training

To learn the generator *G*, we employ adversarial training where a discriminator function *D* parameterized by Θ_*D*_ is trained to distinguish between *h* = *F*(*x*), with *x* ~ **X**^s^, and *ĥ* = *G*(*x*, *F*(*x*)), with *x* ~ **X**^t^ (Figure 6). *D* is a NN-based classifier which estimates the probability of the input latent code coming from the source batch. Therefore, an ideal discriminator would produce *D*(*h*) ≈ 1 and *D*(*ĥ*) ≈ 0. Simultaneously, *G* is optimized to fool the discriminator into misinterpreting *ĥ* as *h*. The two models *G* and *D* are jointly trained to learn parameters Θ_*G*_ and Θ_*D*_ by parallely minimising the corresponding generator and discriminator losses 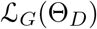 and 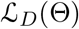 given as,

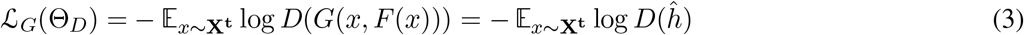

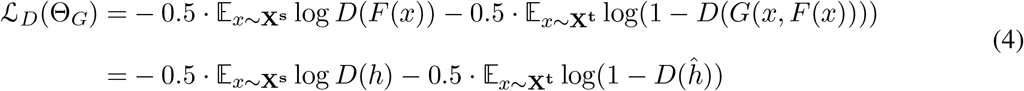

**Figure 6:**
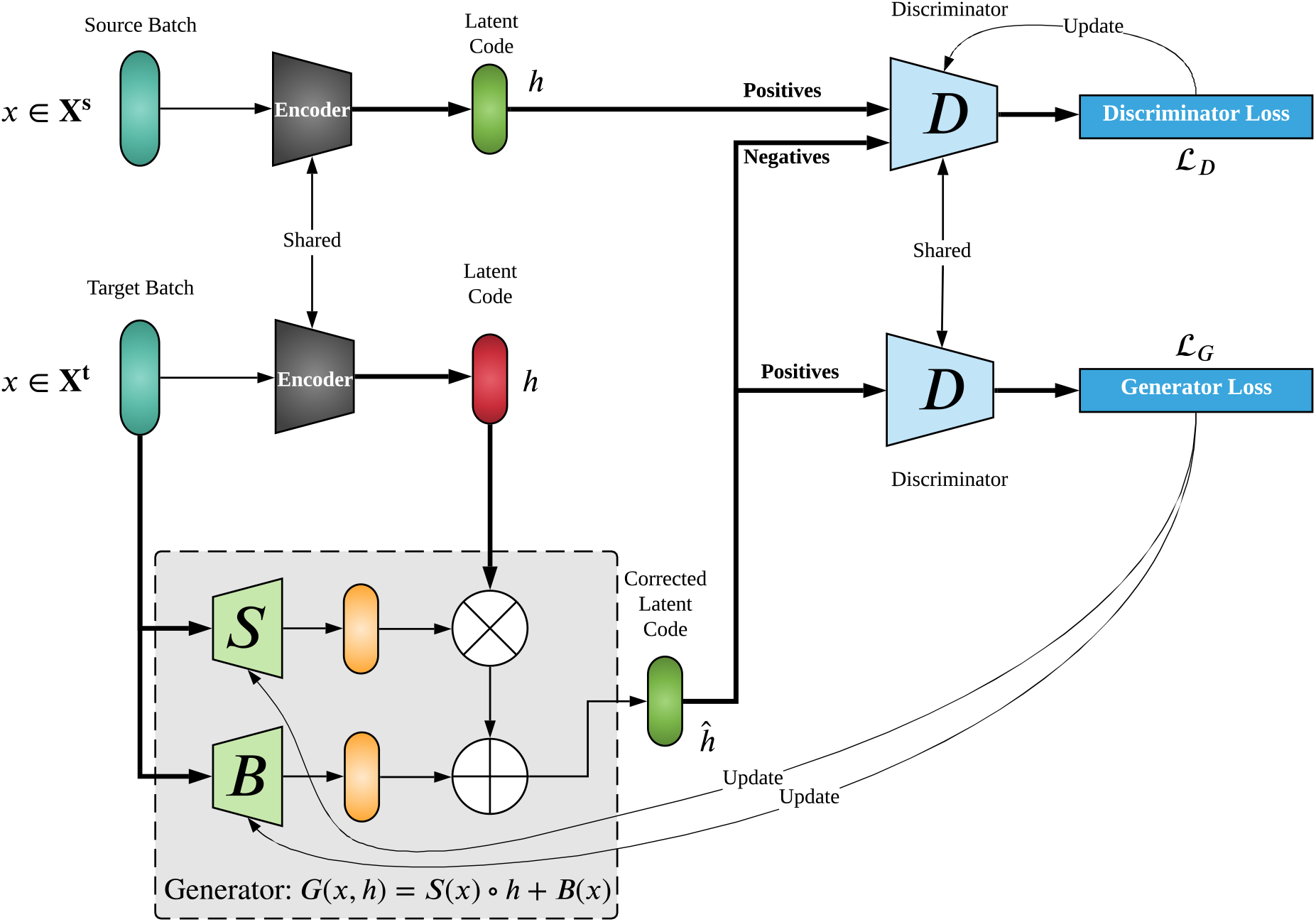
Asymmetric integration used by JIND: A generator scales and shifts the latent code of the target batch, via the NN-based models *S* and *B*, to make it indistinguishable from the latent code of the source batch. To find the optimal parameters, adversarial training is used, in which the generator and a discriminator are jointly trained to minimize their respective losses. The goal of the discriminator is to detect whether the latent code was produced with the source batch (positive examples) or from the target batch (negative examples).

While minimizing 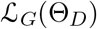 and 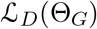, Θ_*D*_ and Θ_*G*_ are kept fixed, respectively. Note that the generator’s objective function only depends on *ĥ*, whereas the discriminator’s objective function depends on both *ĥ* and *h* (Figure 6). JIND’s asymmetric integration requires solving a competitive optimization problem in which two networks with opposite objectives are trained against each other and a saddle point needs to be achieved. This makes the optimization of the generator and discriminator pair challenging. To make the optimization more stable, we regularize the generator’s objective by forcing *S*(*x*) to be approximately **1**. More precisely, we update the generator’s loss function as

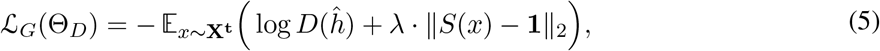

where λ is a hyperparameter (set to 0.001 by default). The goal of this regularization is to prevent the transformation on the target batch from being very complex. Notice that, a more complex generator could learn an incorrect mapping by aligning different cell-types together, which would translate into incorrect predictions for those cells.

#### Network architectures

The generator is composed of *S*(*x*) and *B*(*x*), both with the same network architecture consisting of two fully connected layers containing 512 neurons each (with ReLU activation function) and a fully connected layer at the end which produces a 256-dimensional output. *S*(*x*) and *B*(*x*) each have their own network parameters (contained in Θ_*G*_), and both take as input the cell’s gene expression vector *x* (of dimension 5000 by default).

The network for the discriminator consists of three fully connected layers containing 512, 256 and 256 neurons, respectively (each with Leaky ReLU^44^ activation function), followed by a fully connected layer with one neuron which produces a one-dimensional output between zero and one through sigmoid activation function. The input to the discriminator is a 256-dimensional vector representing the latent code.

#### Training details

We use both the source batch **X**^s^ and the target batch **X**^t^ for training, with no cell annotations. The generator’s parameters Θ_*G*_ are initialized such that initially *G* is an identity mapping, with *h* = *ĥ*, i.e., *S*(*x*) ≈ **1** and *B*(*x*) ≈ **0**. For the generator model, we use RMSProp^45^ optimizer with a learning rate of 0.0001 and a weight decay of 0.01. Similarly, for the discriminator model, we use RMSProp optimizer with a learning rate of 0.0001 and a weight decay of 1e-6. Note that weight decay is equivalent to penalty on the magnitude of the network parameters, which is required to prevent over-fitting. Both architectures use a mini-batch size of 512 samples for the optimization via gradient descent. Since the task of the generator is only to learn the residual, i.e., modify the latent code, for every iteration of the generator the discriminator undergoes two training iterations.

#### Final prediction model

After the asymmetric integration is performed, the final prediction model is as follows. First, the encoder subnetwork takes as input the cell’s gene expression vector *x* and produces the latent code *h*. Both *x* and *h* are then input to the generator, which produces the corrected latent code *ĥ*. Finally, the corrected latent code *ĥ* goes through ReLU activation, and then to the classifier subnetwork, which produces the final probabilities of the cell belonging to each of the considered *K* cell-types (Figure 1c). JIND labels the cell with the cell-type having highest probability, or with an “unassigned” label if the threshold constraint is not satisfied.

### JIND+

JIND+ is an extension of JIND that uses self-training to improve its performance. Self-training involves the use of unlabelled data, in our case the target batch with unknown cell-type annotations, to improve the classification model by using the predictions as pseudo labels. The distribution of the unlabelled data might differ slightly from that of the training data, and hence self-training becomes an efficient way of transferring classifiers across domains^22, 23^. In our case, even after asymmetric integration in the latent space, the distribution of the modified latent code for the target batch may still differ from that of the latent code for the source batch. Therefore, in JIND+ we use the confident predictions made on the target batch to further fine-tune (train) the encoder and the classifier subnetworks. We minimize weighted categorical cross entropy, and reuse the weights determined using source batch cell composition (Eq. 1). To identify the confident predictions, we calculate thresholds for each cell-type as outlined in subsection (filtering) using the validation dataset (from the source batch) with an outlier fraction *θ* = 0.3. Then, only the cells with predictions above the corresponding threshold are used for the fine tuning, using the predicted cell-type as the label. Note that JIND+ performs self-training after training the generator and discriminator networks, after which their parameters remain unchanged. Only the parameters of the prediction model are modified.

#### Training Details

We use Adam optimizer^42^ with a learning rate of 0.0001 for 10 epochs and a mini-batch size of 32. The rest of the hyperparameters are the same as the ones used for training the prediction model. To obtain training and validation datasets, we choose 70% of the confident predictions for training and the rest for validation. Then, we save the parameters across all epochs with the highest accuracy on the validation set (as done in the parameter selection for the prediction model).

## Notes

### Competing Interest Statement

The authors have declared no competing interest.

