## Supplementary Data for "JIND: Joint Integration and Discrimination for Automated Single-Cell Annotation"

### Contents

### List of Tables

S1 Comparing performances of JIND and JIND+ when the source and target batches are integrated with Seurat (Integrated) versus when Seurat integration is not performed (Batched). *raw* is the inital accuracy of the classifier, *rej* is the percentage of cells rejected by the classifier and *eff* is the effective accuracy after rejecting unconfident predictions. Best raw accuracy rates among the two cases (batched or integrated) are **boldfaced**. . . . . . . . . . . 4

S2 Impact of choosing different number of genes for training the prediciton model. NN is the neural network based prediction model used by JIND and doesn't perform any batch alignment. JIND performs adversarial training to minimize distributional mismatch between source and target batches. JIND+ further fine-tunes assuming the confident predictions are correct on the target batch. *raw* is the inital accuracy of the classifier, *rej* is the percentage of cells rejected by the classifier and *eff* is the effective accuracy after rejecting unconfident predictions. . . . . . . . . . . . . . . . . . . . . . . . . . . . . . . . . . . . . . . . . . . 4

S3 Comparison position of ReLU activation w.r.t encoder. Case 1: When ReLU activation is a part of the classifier subnetwork and is applied on the latent code produced by the encoder subnetwork. Case 2: When ReLU activation is a part of encoder subnetwork applied before at the end of encoder to produce the latent code. JIND performs adversarial training to minimize distributional mismatch between source and target batches. JIND+ further fine-tunes assuming the confident predictions are correct on the target batch. *raw* is the inital accuracy of the classifier, *rej* is the percentage of cells rejected by the classifier and *eff* is the effective accuracy after rejecting unconfident predictions. . . . . . . . . . . . . . . . . . . . . 5

### Additional results

- Figure S1 shows different gene expression patterns across the two groups, G1: *Ductal* cells predicted as *Ductal* and , G2: *Ductal* cells predicted as *Acinar*, allowing differentiation between the two populations of cells. On the contrary, when DE analysis was performed on the groups, G1: randomly chosen subset of *Ductal* cells and , G2: remaining *Ductal* cells, (Figure S3) we neither observe meaningful clustering nor descriptive gene expression patterns necessary for differentiation. We can see the same results on the PBMC dataset on Figure S2 and Figure S4

- Table S1 shows the comparison between two cases, Batched: when JIND and JIND+ are evaluated on datasets containing batch effects versus, Integrated: when JIND and JIND+ are run after Seurat integration on the same datasets.

| Datasets | | PBMC 10x_v3-10x_v5 | | Pancreas Bar16-Mu16 | | Pancreas Bar16-Seg16 | |
|---|---|---|---|---|---|---|---|
| | | Batched | Integrated | Batched | Integrated | Batched | Integrated |
| *JIND* | *raw* | **0.971** | 0.968 | **0.958** | 0.946 | **0.987** | 0.946 |
| | *rej* | 0.07 | 0.06 | 0.05 | 0.10 | 0.05 | 0.08 |
| | *eff* | 0.986 | 0.985 | 0.974 | 0.979 | 0.997 | 0.979 |
| *JIND+* | *raw* | **0.974** | 0.971 | 0.959 | **0.961** | **0.992** | 0.961 |
| | *rej* | 0.03 | 0.03 | 0.03 | 0.09 | 0.02 | 0.05 |
| | *eff* | 0.985 | 0.978 | 0.971 | 0.980 | 0.997 | 0.980 |

**Table S1:** Comparing performances of JIND and JIND+ when the source and target batches are integrated with Seurat (Integrated) versus when Seurat integration is not performed (Batched). *raw* is the inital accuracy of the classifier, *rej* is the percentage of cells rejected by the classifier and *eff* is the effective accuracy after rejecting unconfident predictions. Best raw accuracy rates among the two cases (batched or integrated) are **boldfaced**.

| Methods | Datasets | Human-Hemato | | | | Mouse Cortex | | | |
|---|---|---|---|---|---|---|---|---|---|
| | Metrics/#Genes | 1000 | 3000 | 5000 | 7000 | 1000 | 3000 | 5000 | 7000 |
| NN | *raw* | 0.923 | 0.938 | 0.928 | 0.933 | 0.975 | 0.976 | 0.982 | 0.981 |
| | *rej* | 0.06 | 0.06 | 0.06 | 0.06 | 0.05 | 0.05 | 0.05 | 0.06 |
| | *eff* | 0.945 | 0.957 | 0.950 | 0.954 | 0.988 | 0.990 | 0.991 | 0.997 |
| JIND | *raw* | 0.922 | 0.939 | 0.927 | 0.931 | 0.976 | 0.970 | 0.982 | 0.981 |
| | *rej* | 0.07 | 0.06 | 0.06 | 0.06 | 0.06 | 0.03 | 0.05 | 0.06 |
| | *eff* | 0.944 | 0.957 | 0.948 | 0.953 | 0.989 | 0.982 | 0.991 | 0.996 |
| JIND+ | *raw* | 0.919 | 0.931 | 0.927 | 0.932 | 0.976 | 0.970 | 0.976 | 0.978 |
| | *rej* | 0.04 | 0.04 | 0.04 | 0.04 | 0.02 | 0.03 | 0.04 | 0.04 |
| | *eff* | 0.933 | 0.945 | 0.943 | 0.948 | 0.985 | 0.950 | 0.984 | 0.990 |

**Table S2:** Impact of choosing different number of genes for training the prediciton model. NN is the neural network based prediction model used by JIND and doesn't perform any batch alignment. JIND performs adversarial training to minimize distributional mismatch between source and target batches. JIND+ further fine-tunes assuming the confident predictions are correct on the target batch. *raw* is the inital accuracy of the classifier, *rej* is the percentage of cells rejected by the classifier and *eff* is the effective accuracy after rejecting unconfident predictions.

| Methods | Datasets | PBMC 10x_v3-10x_v5 | | Pancreas Bar16-Mur16 | | Pancreas Bar16-Seg16 | |
|---|---|---|---|---|---|---|---|
| | Metrics | Case 1 | Case 2 | Case 1 | Case 2 | Case 1 | Case 2 |
| JIND | *raw* | 0.971 | 0.972 | 0.958 | 0.881 | 0.987 | 0.86 |
| | *rej* | 0.07 | 0.07 | 0.05 | 0.39 | 0.05 | 0.44 |
| | *eff* | 0.986 | 0.988 | 0.974 | 0.987 | 0.997 | 0.9948 |
| JIND+ | *raw* | 0.974 | 0.974 | 0.959 | 0.8805 | 0.992 | 0.901 |
| | *rej* | 0.03 | 0.03 | 0.03 | 0.27 | 0.02 | 0.23 |
| | *eff* | 0.985 | 0.984 | 0.971 | 0.979 | 0.997 | 0.984 |

**Table S3:** Comparison position of ReLU activation w.r.t encoder. Case 1: When ReLU activation is a part of the classifier subnetwork and is applied on the latent code produced by the encoder subnetwork. Case 2: When ReLU activation is a part of encoder subnetwork applied before at the end of encoder to produce the latent code. JIND performs adversarial training to minimize distributional mismatch between source and target batches. JIND+ further fine-tunes assuming the confident predictions are correct on the target batch. *raw* is the inital accuracy of the classifier, *rej* is the percentage of cells rejected by the classifier and *eff* is the effective accuracy after rejecting unconfident predictions.

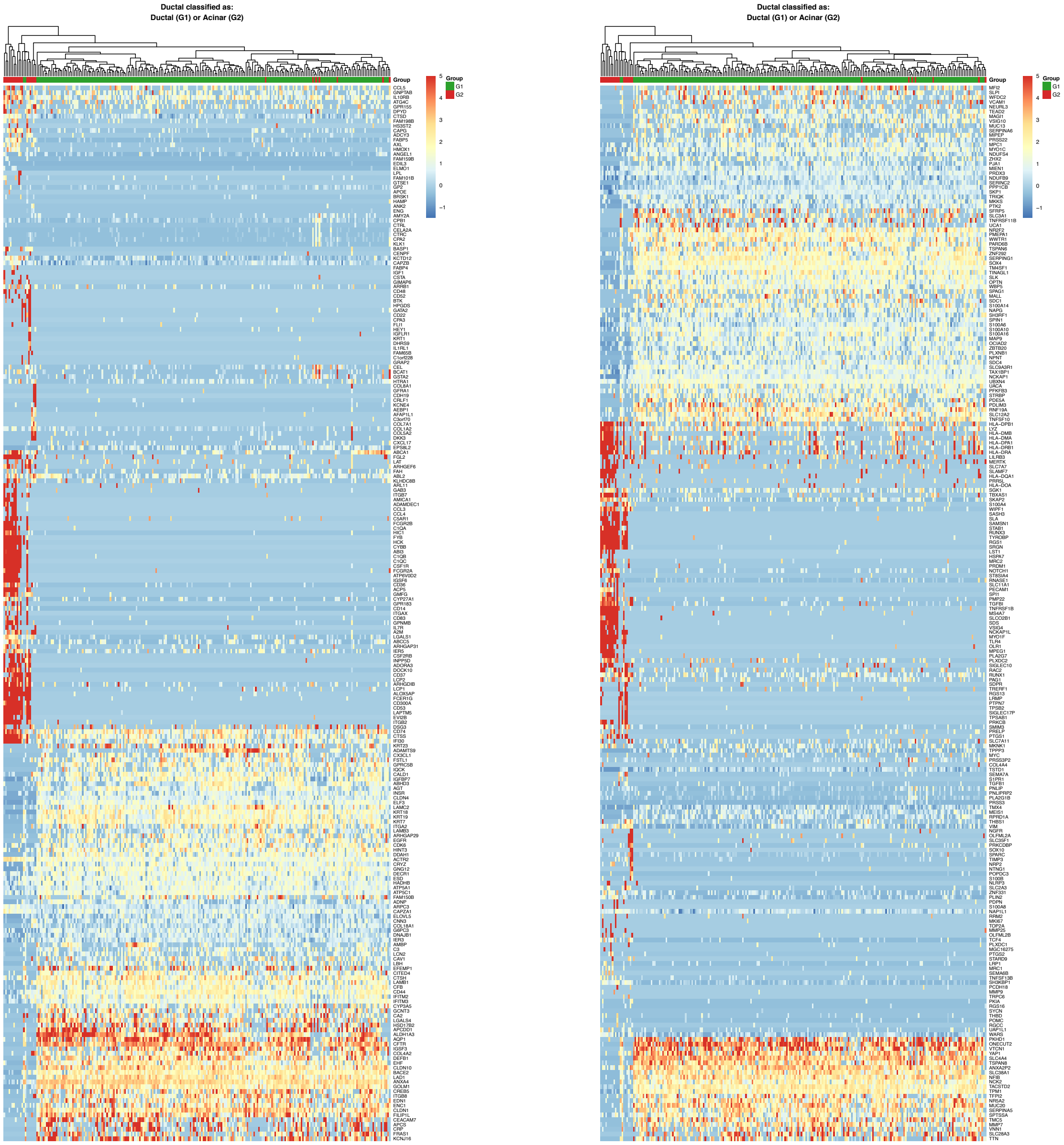

**(a)** First half of the DEG

**(b)** Second half of the DEG

**Figure S1:** Heatmap for all differentially expressed genes between two groups on *PBMC* 10x_v5 dataset. *Monocytes FCGR3A* cells predicted by JIND+ as: *Monocytes FCGR3A* cells (G1) or *Monocytes CD14* cells (G2). The clustering of the cells is calculated with the entire set of genes. The results of this DE analysis can be found on the Supplementary Excel File.

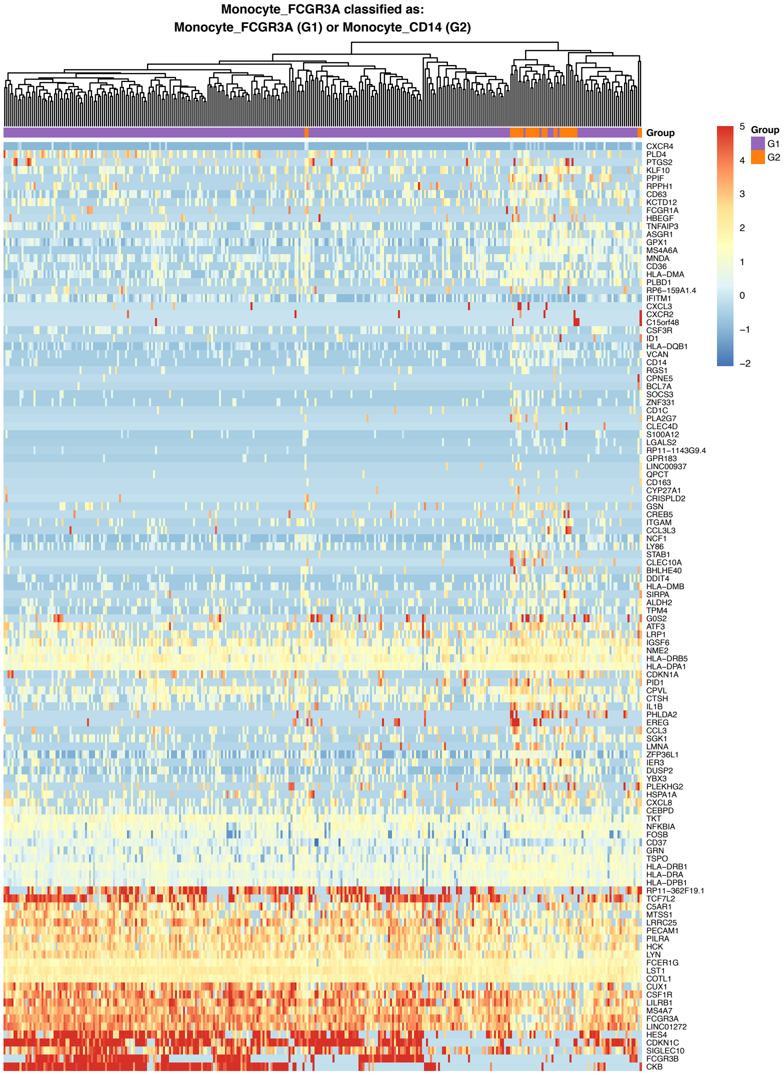

**Figure S2:** Heatmap for all differentially expressed genes between two groups on *PBMC* 10x_v5 dataset. *Monocytes FCGR3A* cells predicted by JIND+ as: *Monocytes FCGR3A* cells (G1) or *Monocytes CD14* cells (G2). The results of this DE analysis can be found on the Supplementary Excel File.

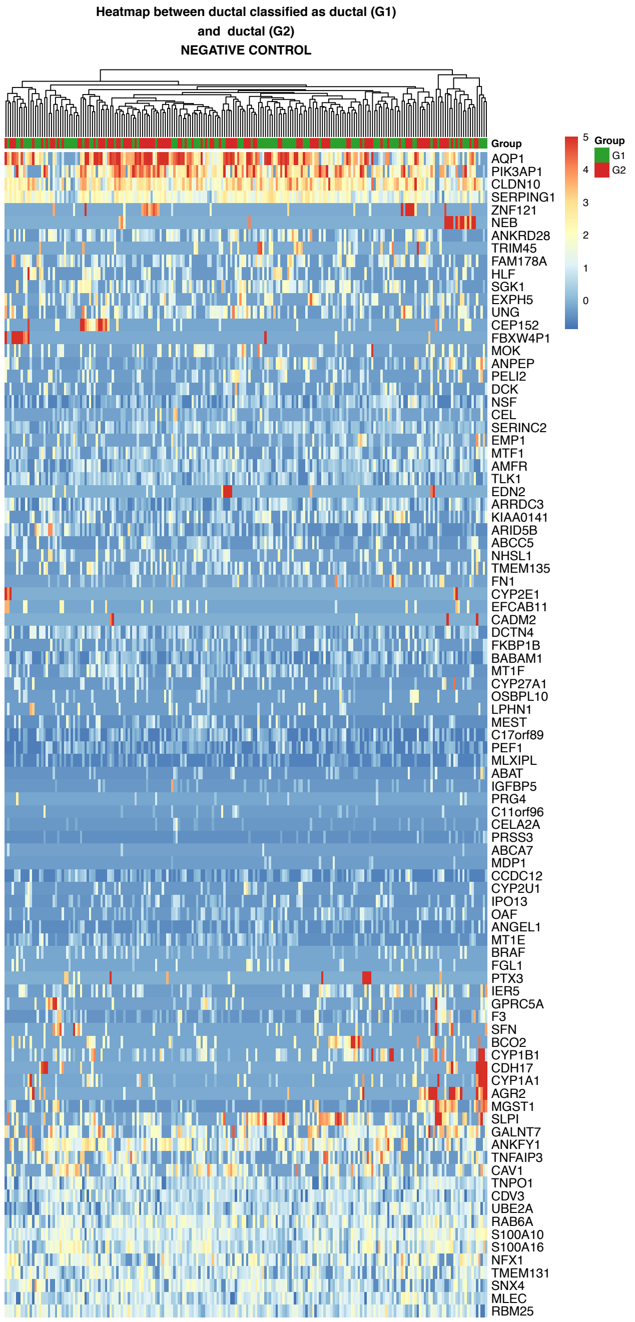

**Figure S3:** Heatmap for all differentially expressed genes among two randomly chosen groups of *Acinar* cells present in *Pancreas* Mur16 dataset. The results of this DE analysis can be found on the Supplementary Excel File.

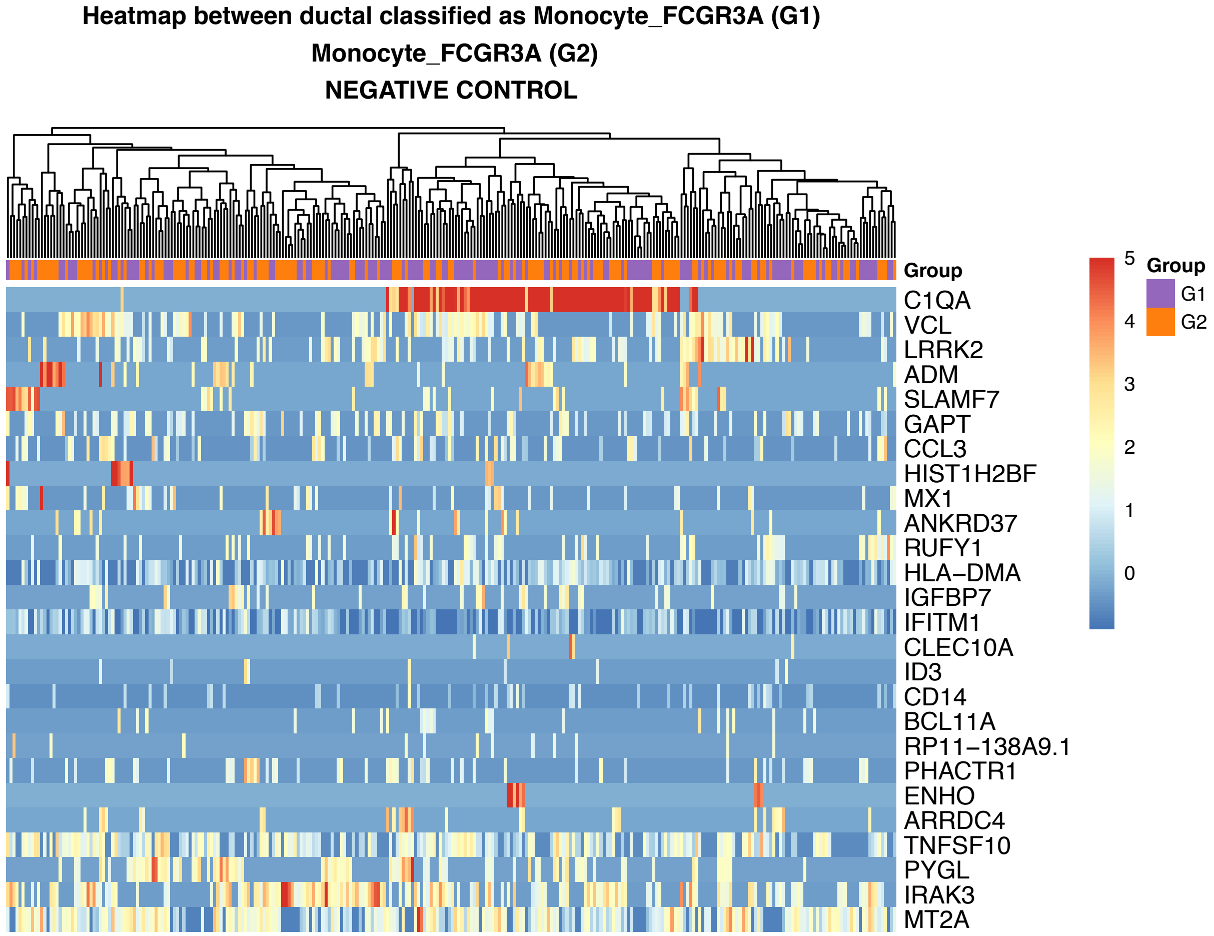

**Figure S4:** Heatmap for all differentially expressed genes among two randomly chosen *Monocyte FCGR3A* cells present in *PBMC* 10x_v5 dataset. The results of this DE analysis can be found on the Supplementary Excel File.

### Neural Network based Prediction Model

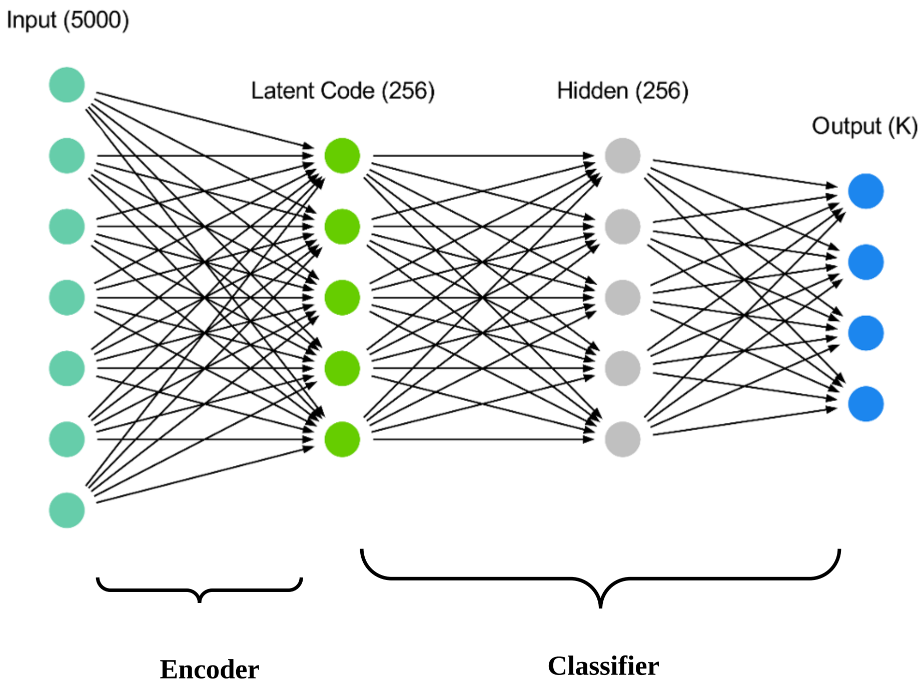

**Figure S5:** NN-based prediction model employed by JIND for cell identification. The network consists of two subnetworks, an encoder and a classifier, which are jointly trained. The input to the model is a vector (of dimension 5000 by default) containing the gene expression data for a cell, and the output consists of a probability vector indicating the likelihood of the cell belonging to each of the $K$ classes.
